# An approach to stress assessment during hunting: Cortisol levels of wild boar (*Sus scrofa*) during drive hunts

**DOI:** 10.1101/2020.03.11.987628

**Authors:** Justine Güldenpfennig, Marion Schmicke, Martina Hoedemaker, Ursula Siebert, Oliver Keuling

**Affiliations:** Institute for Terrestrial and Aquatic Wildlife Research, University of Veterinary Medicine Hannover, Foundation, Bischofsholer Damm 15, 30173 Hannover, Germany; Institute of Agricultural and Food Sciences, Animal Health Management, Martin Luther University Halle-Wittenberg, Theodor-Lieser-Straße 11, 06120 Halle, Germany; Clinic for Cattle, University of Veterinary Medicine Hannover, Foundation, Bischofsholer Damm 15, 30173 Hannover, Germany

**Keywords:** animal welfare, cortisol, game management, hunting, stress, wild boar

## Abstract

Only little is known about the effect of stress (both short-term and long-term) on wildlife species. To get an idea of stress in wildlife, we investigated the cortisol level of wild boar during drive hunts in Lower Saxony, Germany. Cortisol as one of the main stress hormone in mammals is considered to have negative impacts on the animal’s well-being if expressed excessively (repeatedly over a longer period). We analysed serum cortisol levels of 115 samples using a radioimmunoassay and compared sampling month, hunting grounds, age classes and sexes, as well as possible correlations between cortisol level and weight and pregnancy status of female wild boar. We found that cortisol levels during these drive hunts exhibit wide variation. The mean cortisol level was 411.16 nmol/L with levels ranging from 30.60 nmol/L (minimum) to 1,457.92 nmol/L (maximum). Comparing age groups and sexes, we found significant differences between the sexes, with females having a higher cortisol levels than males. After grouping age groups and sexes together, we also found significant differences based on the age-sex group. We found no correlation between cortisol levels and weight, but significantly higher cortisol levels in pregnant females compared to non-pregnant females. No differences were found between sampling months and locations, respectively. These results show the impact of drive hunts on stress in wild boar; nevertheless, this impact of drive hunts as performed in most parts of Central Europe seems to be not as high as imagined. Still, we need more information about cortisol levels and stress in (hunted) wildlife species.

## Introduction

With stress research in humans and domestic animals is well established, little is known about possible stress in wildlife. However, interest in stress research based on animal welfare is growing [1]. Causes for stress in wildlife could be deforestation, habitat fragmentation, insufficient food and water resources as well as human disturbances [2,3]. During stress, the animal’s behaviour is altered to enhance attention, increase cardiac output, respiration, and catabolism, as well as to divert blood flow to provide full perfusion of the brain, heart and muscles [4]. While glucocorticoids are beneficial for short-term survival, enduring/increased release (like during chronic stress) can lead to metabolic, immune and physiological dysfunction [5]. Cumulative occurrence of stressors could lead to changes in an animal’s well-being and to changes in social and disease networks [6,7]. It was found that chronic stress in domestic pigs leads to altered cortisol levels, reduced growth and reduced play-behaviour [8]. High cortisol levels are also linked to many behavioural, physiological, nutritional diseases and disorders [3], as well as to obesity and diabetes caused by increasing plasma glucose concentrations due to cortisol [9]. The relationship between wildlife and disease as well as host-parasite equilibrium can influence animal populations and lead to loss of biodiversity [3]. Therefore, a better understanding of stress in wildlife seems to be necessary. The wild boar *Sus scrofa* (Linnaeus, 1758) is nowadays distributed across almost all continents and shows one of the most widely spread native geographic ranges [10–12]. Because of some traits, such as opportunistic feeding and habitat diversion [11], they may not the best model organism for stress research, assuming relatively low stress responses to chronic stress triggers such as food limitation and habitat loss. However, blood samples of wild boar are easy to collect in large numbers due to high density populations in almost every habitat [11] and an all-year hunting season [13]. In addition, behavioural and social structures are well known [11,12], allowing for the investigation of stress differences between sexes, ages and social positions. There are said to be differences between the responses to chronic stress of males and females, as well as different social ranks in pigs [14]. Additionally, knowledge of wild boar stress and stressors may also help to improve domestic pigs’ welfare in pig farming, by supplementing the available stress research specific to domestic pigs.

Stress is a widely used term in our society. Chronic stress, or the repeated occurrence of a stressor, has negative impacts on well-being [15]. Stress could be defined as “the nonspecific response of the body to any demand of change” [16,17]. In general, stress is a change in the psychological, physiological and/or physical features of an organism [3]. One of the main stress hormones in almost all mammals is cortisol [18]. Cortisol is well established in animal welfare research [19,20] and plays a key role in energy release, immune and mental activity, development and growth, as well as in reproductive functions [21,22]. In its important role in stress response, cortisol is sensitive to both physical and emotional stimuli; the release underlies a circadian rhythm [21]. Under physiologic stress, cortisol modulates the immune system and mobilizes energy storages, making more resources available for responding to a certain stressor [22,23]. Repeated stress impulses can lead to an over activation of the Hypothalamus-Pituitary-adrenal (HPA) axis, which could result in elevated cortisol levels and shorter periods of enhanced cortisol with acute stressors or lower basal levels/hypocortisolism (a deficiency of cortisol) [21,24].

Since we do not know much about the mechanisms and impact of stress in wildlife, this work serves as a pilot study to measure stress in wildlife and investigate the effect of drive hunts on the cortisol levels, and therefore on the acute stress, of game species such as the wild boar. We investigated the following questions: (1) What are the cortisol levels of wild boar during drive hunts and (2) are there differences between hunting months, hunting grounds, age groups and sexes, as well as pregnant and non-pregnant wild boar?

## Materials and Methods

### Study area

The study area was located in the eastern part of the federal state of Lower Saxony (Northern Germany) (52.36°N, 10.35°E) [similar to 25,26]. Altitude ranged from 60 to 130 m above sea level with sub continental climates [27]. The average annual precipitation in 2018 was 512 mm; the average annual temperature in 2018 was 10.7° C [28]. The approximately 4,900 km^2^ area is composed of 54.6 % of cultivated fields and 27.6 % forest, with buildings and infrastructure claiming the remaining 17.8 % [for detailed information see 25,26]. In the study area, hunting is usually performed as a single hunt or a drive hunt. Supplementary feeding is not allowed, but baiting as a necessary hunting strategy is [29]. Wild boar hunting has especially intensified due to risk control management associated with African Swine Fever [30].

### Sample collection and laboratory analysis

Using blood as the sample material for cortisol measurement is often linked with difficulties in evaluation because animal handling itself could cause stress itself [1]. Alternatives used to measure glucocorticoids in other studies include saliva, faeces and hair [1,31]. Although salivary cortisol levels have been considered to be a better way to determine stress in animals, cortisol levels measured in saliva must also be treated with caution, as saliva contains only biologically free cortisol which does not respond linearly to a stressful stimulus [32]. In addition, collecting saliva from dead animals remains challenging as blood often contaminates the snouts of hunted wild boar after death, which has to be avoided [31]. Faeces and hair on the other hand are easy to obtain, especially from dead wild boar with a year-round hunting season [13]. Faeces only contain metabolites of cortisol from the past hours to days, based on metabolism [33]. A much longer record of cortisol levels can be measured using hair, which makes it useful for measuring chronic stress [31]. Regardless, we chose blood for stress determination because it was easy to obtain and allowed for a direct insight into the acute stress situation created during drive hunts [31].

We collected blood from dead wild boar after drive hunts in the western part of Lower Saxony, Germany. The blood was collected using 7.5 ml Kabevettes® (Kabe Labortechnik GmbH, Nümbrecht-Elsenroth) during slaughtering, max. 3 h after death. We centrifuged the blood samples (12 min., 4500 rpm), transferred 1 ml serum to 1.5 ml Eppendorf Tubes® (Eppendorf AG, Hamburg) and stored them at −32° C until assaying. The shot animals were categorised in age classes based on their tooth eruption [27]. We classified animals as j = juveniles < 12 months (piglets), y = yearlings 12 - 23 months or a = adults ≥ 24 months, as well as differentiating between the two sexes f = females and m = males [34]. In total, we chose 115 samples for examination in order to have a similar sample size in all age-sex classes. Laboratory analysis was conducted at the endocrinological laboratory of the clinic for cattle of the University of Veterinary Medicine Hannover, Foundation using radioimmunoassay (Cortisol RIA KIT, Beckmann Coulter, Inc., Krefeld) while following the instructions from the manufacturer. Additionally, we sampled uteri and ovaries for the ongoing analysis of the reproductive status of female wild boar [ccording to techniques described in 27,35,36]. With this information, we categorized female wild boar as potentially pregnant (several corpora lutea, but no visible embryos), pregnant (developed foetuses) and non-pregnant (no visible corpora lutea or foetuses).

Our values showed a very high variance with both highly increased, as well as very low cortisol values. We compared the distribution of our values to reference values given by Gentsch et al. (2018), who defined both “normal” and “trauma” cortisol levels. They defined cortisol concentrations of undisturbed animals (animals not being followed by humans or dogs and therefore, shot immediately without suffering, e.g. during single hunts) as normal. Concentrations of cortisol in animals being shot during regular drive hunts, or animals having died after an injury (e.g. due to a traffic accident), were defined as traumatized [37]. With this definition, they set 249.6 ± 36.8 nmol/L as normal and 775.2 ± 64.7 nmol/L as trauma cortisol levels (mean ± standard error) in wild boar.

### Ethical Statement

All described sampling methods were conducted during or after normal legal hunting activities due to the laws of the Federal Republic of Germany and the Federal State of Lower Saxony and all international and/or institutional guidelines for animal handling were followed. No animals were harmed or killed for our sampling. No ethical permit for animal experiments applies or has to be permitted. This study did not use human participants.

### Statistical analysis

All statistics were done with R version 3.5.2 [38]. We tested for significant differences between cortisol levels grouped by sampling month, hunting grounds, state of pregnancy in female wild boar, age group and sex, as well as by age and sex combined. The confidence level was set as α = 0.05. We additionally used the FSA package [39], with p-values adjusted using the Benjamini-Hochberg method, and the ggpubr package [40] for R.

## Results

Viewing our results in clusters comparable to levels found by Gentsch et al. (2018), it is noticeable that only 54 % of our cortisol values could be considered as increased trauma levels (>350 nmol/L), whereas 38 % of our values could be defined as normal cortisol levels (150 – 350 nmol/L). The remaining 8 % are out of range. The mean cortisol level and standard deviation of the total sample is 411.2 ± 242.6 nmol/L; the median is 364.7 nmol/L. Minimum and maximum cortisol levels are 30.6 nmol/L and 1457.9 nmol/L, respectively. The mean cortisol levels of shot animals in October, November, December and January are 497.2 ± 354.2 nmol/L, 404.2 ± 263.4 nmol/L, 383.5 ± 236.7 nmol/L and 412.9 ± 166.2 nmol/L, respectively (Fig. 1). Although our values show small differences, there are no significant differences among months (Kruskal-Wallis chi-squared = 1.8848, df = 3, p-value = 0.5967).

**Figure 1:**
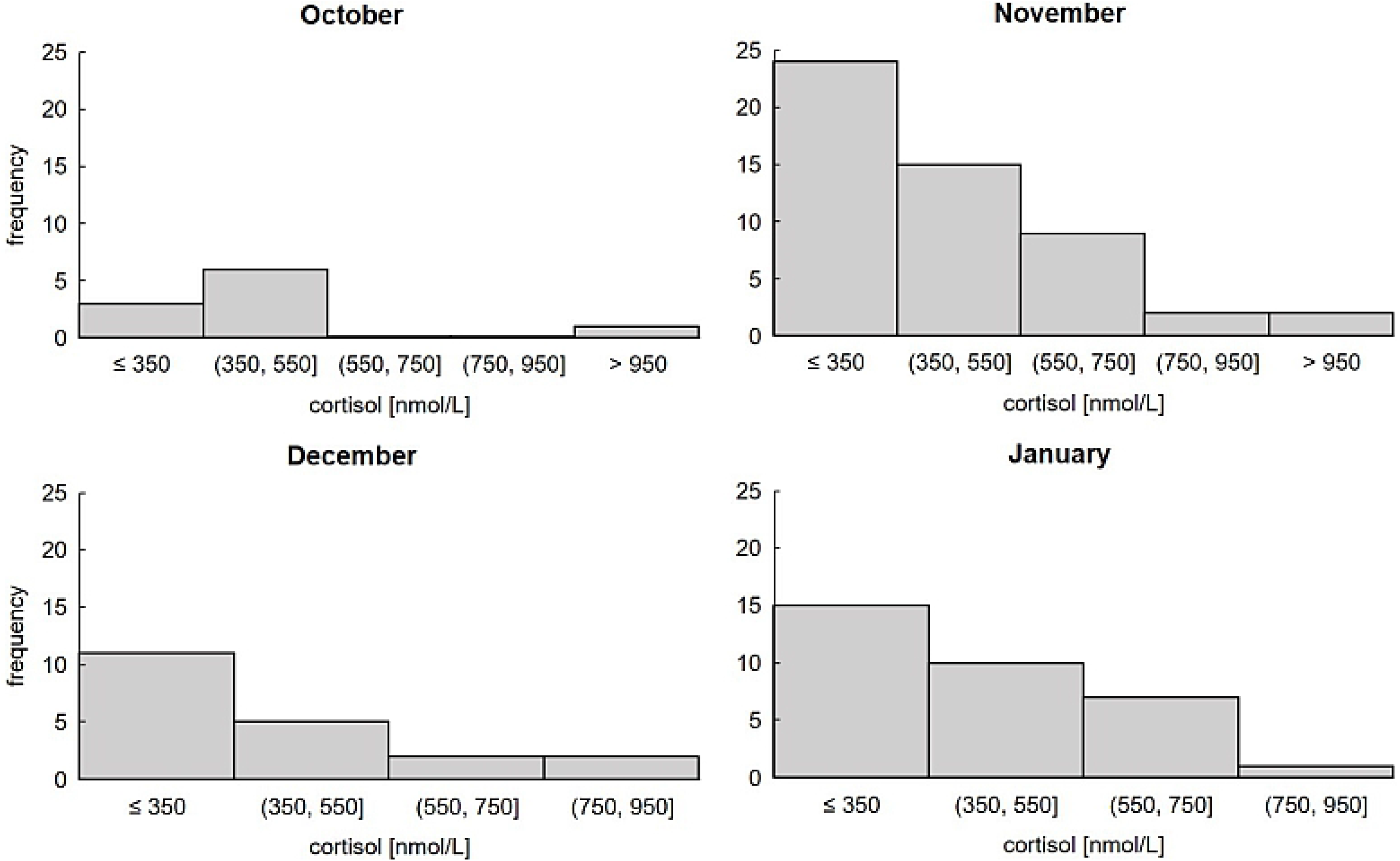
Cortisol levels of wild boar clustered in blocks matching Gentsch et al. 2018. We defined clusters of cortisol levels below 350 nmol/L as normal/basis levels and anything above as increased trauma levels caused by increased stress during drive hunts and compared the hunting months October (n = 10), November (n = 52), December (n = 20) and January (n = 33). There is no difference between hunting months.

The mean cortisol levels show high variation between the different hunting districts (Fig. 2). However, due to the small sample size, there are no significant differences between hunting districts (Kruskal-Wallis chi-squared = 35.701, df = 27, p-value = 0.122).

**Figure 2:**
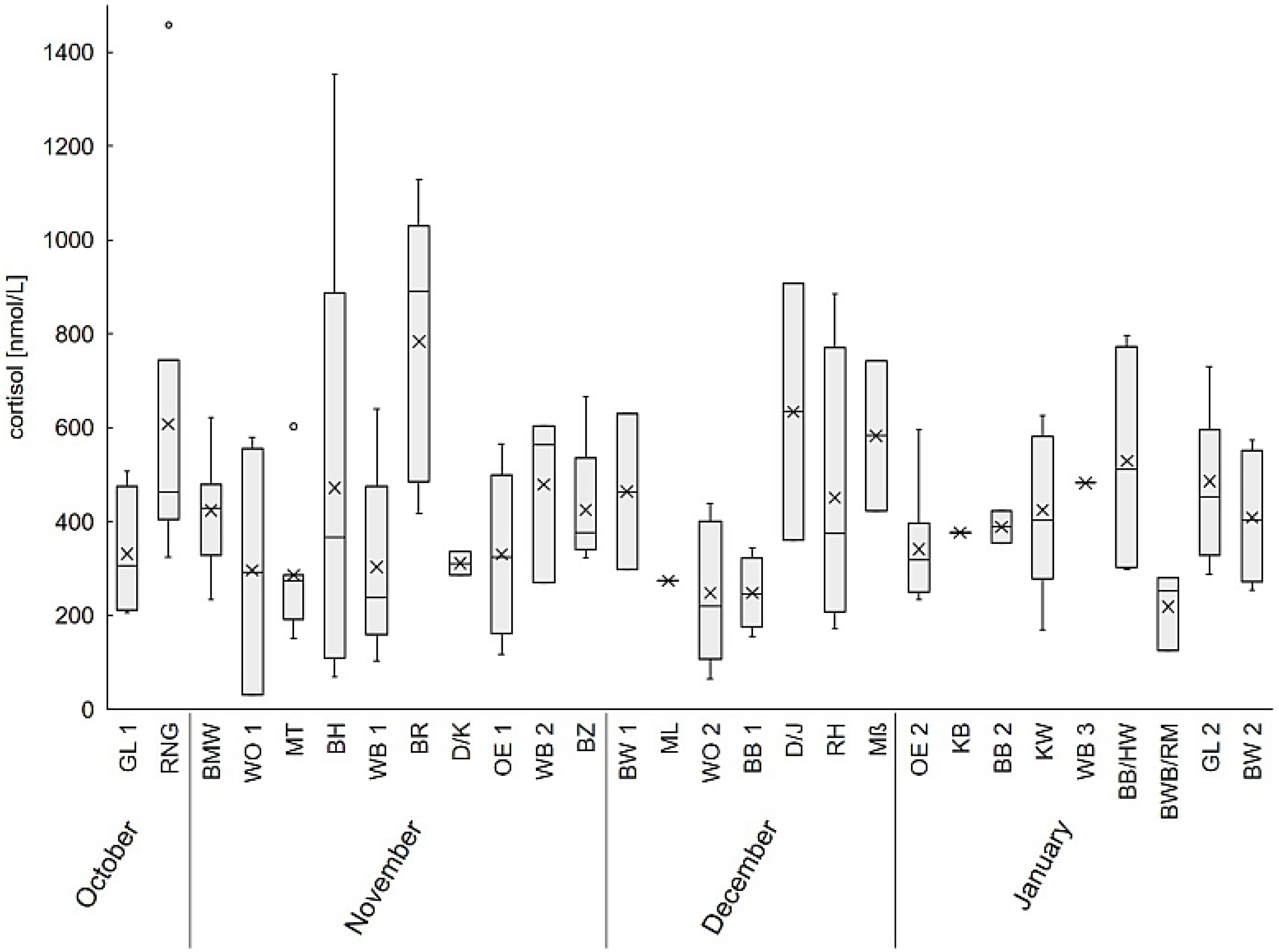
Cortisol levels of wild boar compared between all evaluated hunting grounds. The sampling locations are sorted chronologically from October until January. The abbreviations serve as orientation for the different hunting grounds, which will not named further because of protection of privacy.

We additionally tested for correlation between cortisol levels and dressed weight (weight after gutting), but did not find any correlation (Pearson’s product-moment correlation: t = −0.63456, df = 96, p-value = 0.5272, cor = −0.06462947; Fig. 3).

**Figure 3:**
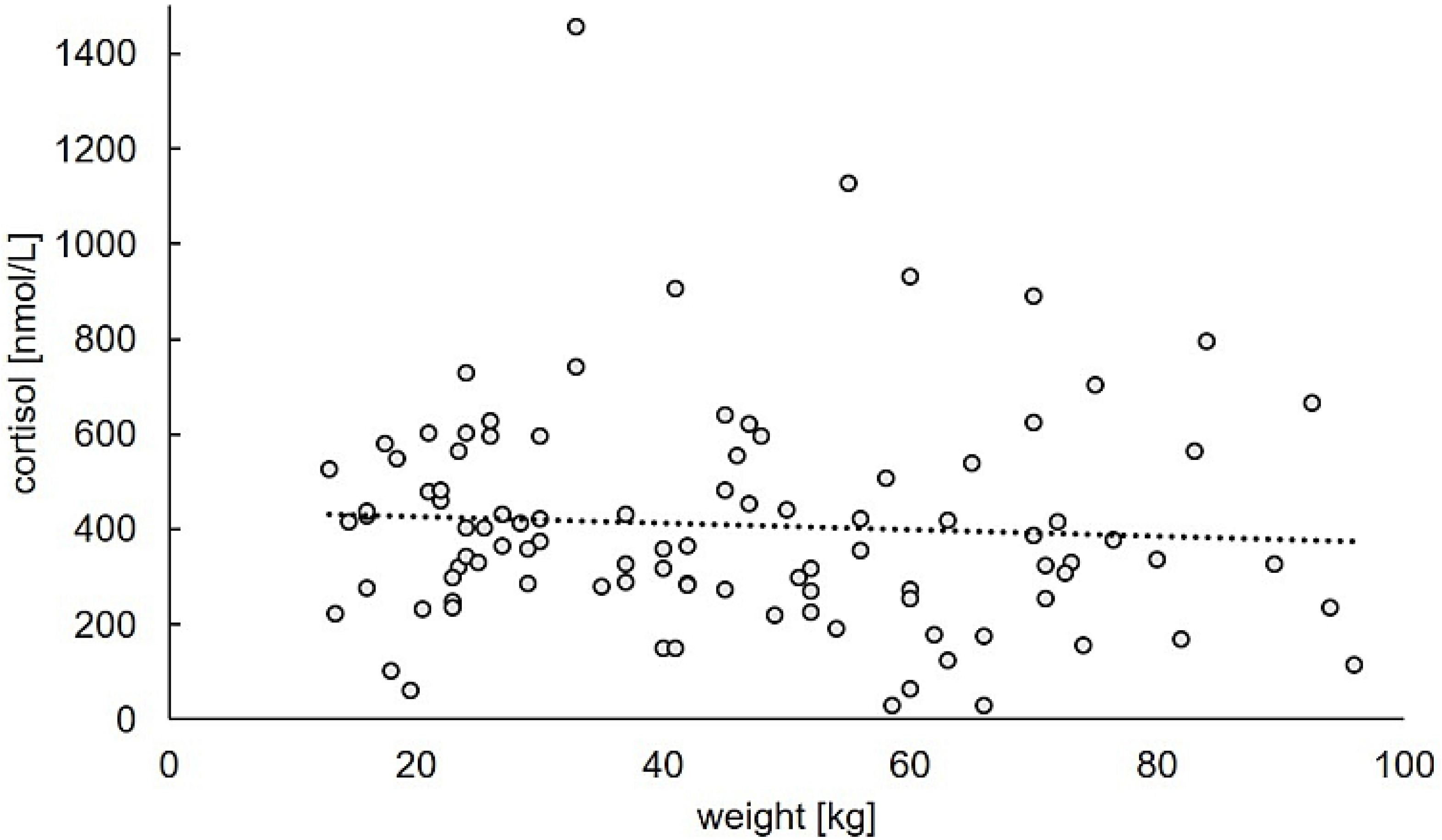
Correlation between cortisol levels and weight of wild boar. The values are highly variable and no correlation could be found. Regression line: y = − 0.6801x + 440.79; R^2^ = 0.0042.

Female wild boar have a significantly higher cortisol levels during drive hunts than do male wild boar (469.65 ± 241.99 nmol/L compared to 353.67 ± 230.97 nmol/L for female and male, respectively; Kruskal-Wallis chi-squared = 11.491, df = 1, p = 6.993 × 10^−4^). Comparing age groups, there are no significant differences (435.04 ± 233.52, 369.63 ± 273.89 and 418.19 ± 221.41 nmol/L for age group 0, 1 and 2, respectively; Kruskal-Wallis chi-squared = 4.326, df = 2, p-value = 0.115). Grouping sex and age classes, the lowest cortisol levels are found in male yearlings, whereas the highest value can be found in female adults (Fig. 4).

**Figure 4:**
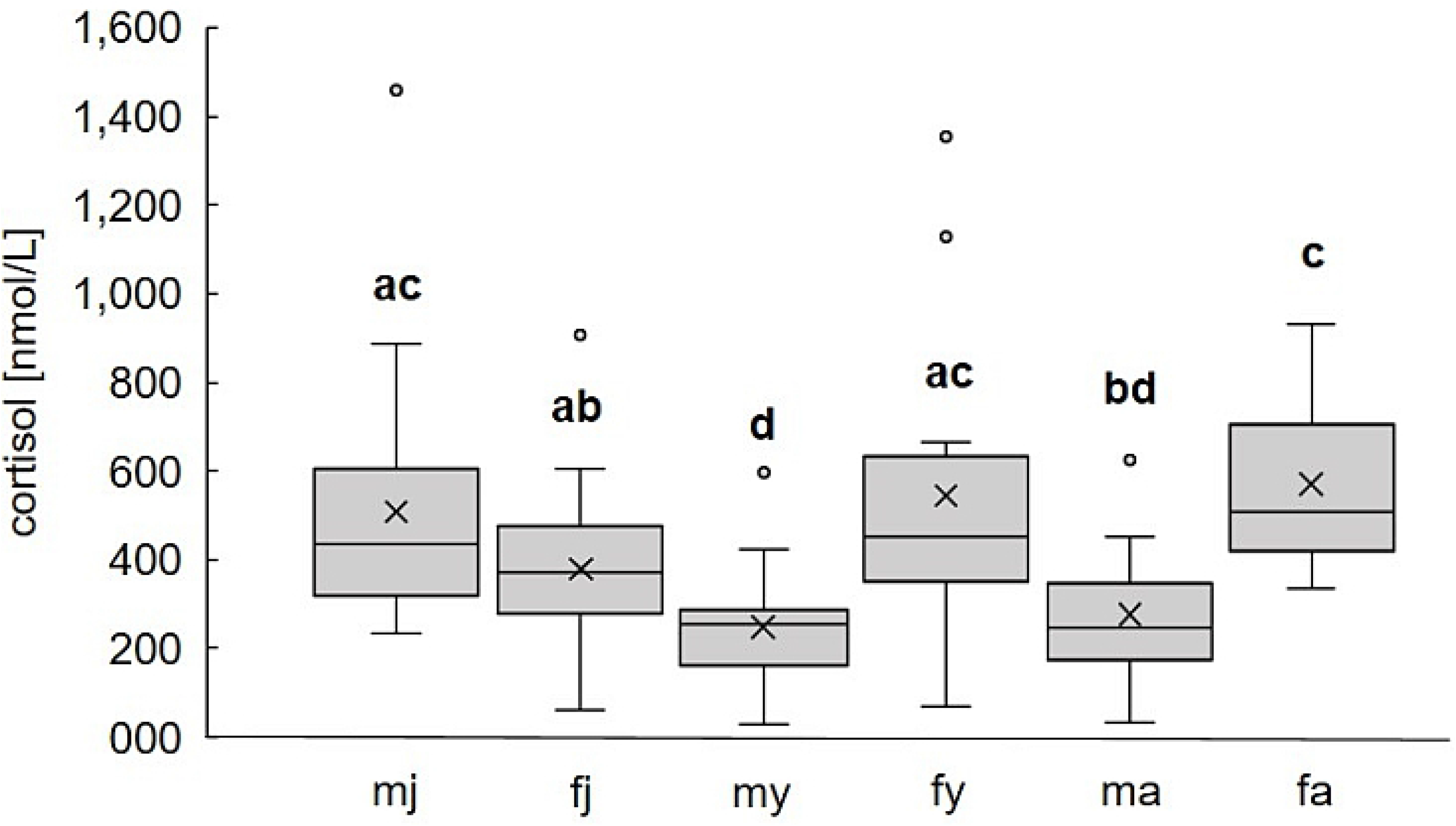
Differences of cortisol levels of wild boar between each age class and sex. Female wild boar (f, n = 57) in general showed a higher cortisol concentration than male (m, n = 58). The age and gender groups combined show various differences, with adult females (fa) having the highest and male yearlings (my, blue) having the lowest cortisol levels. Different letters indicate significant difference (e.g. significant higher cortisol levels in female adults compared to male adults, male yearlings and female juvenile).

Non-pregnant and potentially pregnant wild boar show similar cortisol levels. After separating the cortisol values of definitely pregnant females, these animals were found to exhibit significantly higher cortisol levels, as compared to non-pregnant females (Kruskal-Wallis chi-squared = 4.1683, df = 1, p-value = 0.0412; Fig. 5). The number of foetuses does not have an influence on cortisol levels (Kruskal-Wallis chi-squared = 11.06, df = 7, p-value = 0.136).

**Figure 5:**
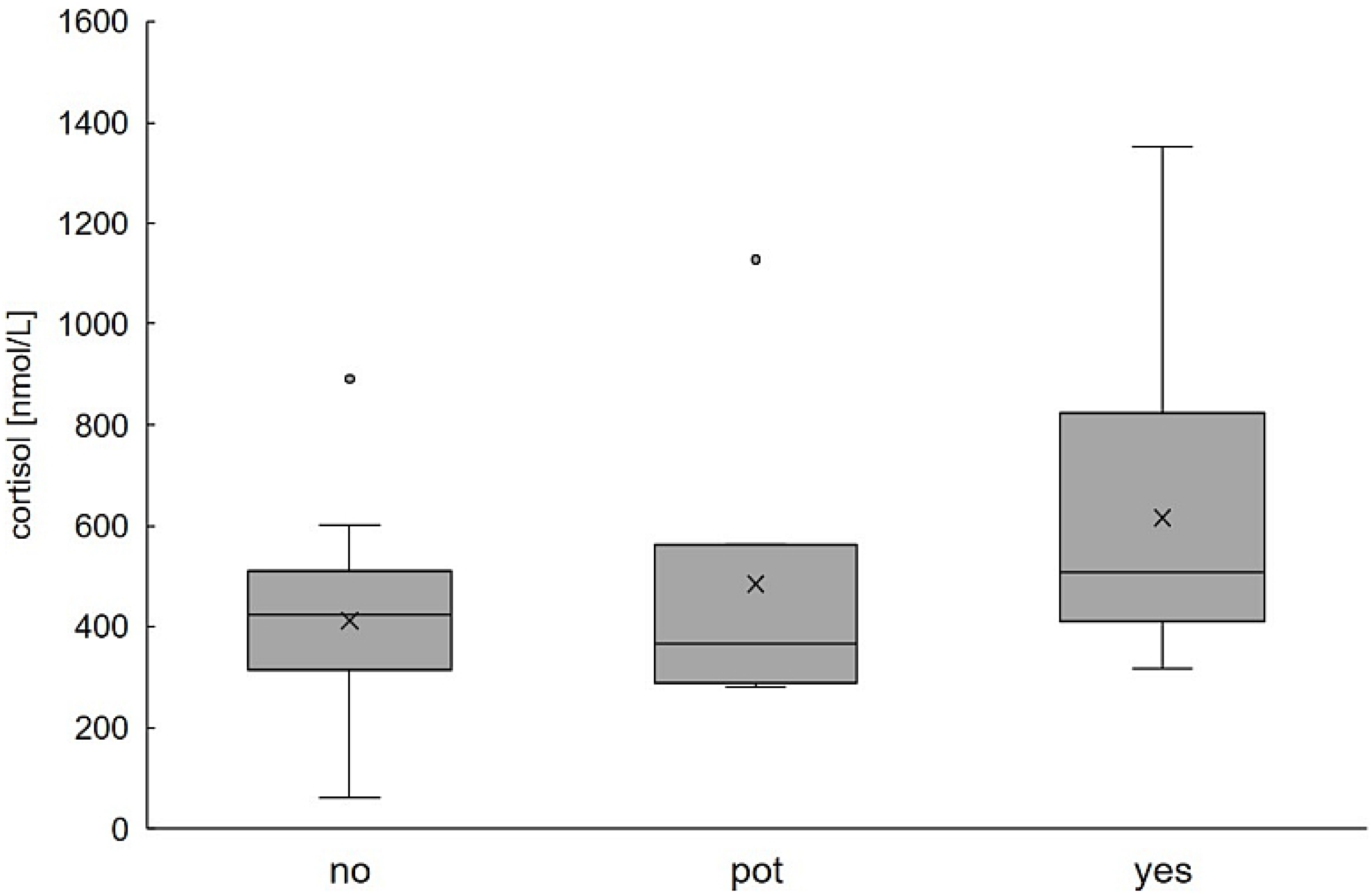
Concentration of cortisol levels of non-pregnant (no, n = 26), potentially pregnant (pot, measurable number of Corpora lutea, n = 7) and definitely pregnant (yes, development of foetuses, n = 14) wild boar. Although there is a noticeable increased cortisol level in definitely pregnant wild boar, there are no significant differences among groups.

## Discussion

We assumed that cortisol levels would naturally increase during drive hunts, as cortisol is one of the main hormones serving to increase energy release during a stress event [21]. Our results show a very high variance with both highly increased, but also very low, cortisol values. Given the fact that we sampled after drive hunts, our results indeed show a high percentage of trauma cortisol levels (54 %), but not as high as was expected. Therefore, while the effect of hunting stress is present, the way we conduct drive hunts in Lower Saxony and most parts of Central Europe seems to be less stressful for wild boar than we expected. There is almost no difference between hunting months and hunting districts. This shows that there is also no long-term effect of drive hunts on wild boar, as cortisol levels did not increase even after repeated drive hunts in the same area over the span of one or two months. Still, we must keep in mind that we do not know the exact condition of the wild boar before death. Differences in individual behaviour, e.g. the duration of them being chased, could determine their cortisol levels.

The given differences between male and female wild boar, with females having an overall higher cortisol level compared to males (469.65 ± 241.99 nmol/L compared to 353.67 ± 230.97 nmol/L, respectively), were very interesting to note. Female wild boar form social groups consisting mostly of mothers with their offspring, and are therefore considered to be matrilineal [11,12,41,42]. Male wild boar leave the group when reaching puberty [43]. Despite the benefits of association in female wild boar, costs like resource competition [44] could lead to specific social pressure and might result in higher cortisol concentrations in females, as well as in group living male piglets (juvenile male). Differences in cortisol levels between sexes, as well as between social ranks, were also found in other species [45,46], though the results seem to be contentious, depending on what assay was used. For instance, it was shown that growing pigs at the age of 4, 8 and 12 weeks exhibit no gender-based cortisol level differences [47], similar to pigs until the weight of 104 ± 7 kg live body mass [48]. This supports our findings that there are no differences in the cortisol levels of female and male juvenile wild boar (Fig. 4). Other studies found significantly higher cortisol levels in male Yucatan minipigs, as compared to females [14], and in barrows compared to gilts [49]. In contrast to that, higher cortisol levels in females than in males (similar to our results) were found in rodents [50], as well as in domestic pigs [9] and in humans before and after a rowing ergometer competition [51]. Rats also show a greater cortisol increase in females after a stress event than do males [52]. In humans, the gender-based differences seem to be more complex [45]. One study revealed no differences in basal cortisol levels between men and women, a greater cortisol increase in men due to math and verbal challenges, and a greater cortisol increase in women in response to social rejection challenges [53].

A correlation between cortisol levels and weight could not be found (Fig. 3), regardless of age and gender related differences in weight. This is in contrast to findings showing relationships between cortisol levels and mass-specific metabolic rate, linked to body mass, in other mammals [15].

Another explanation for the differences between sexes, other than the social structure of wild boar, is the general divergence of male and female wild boar based on metabolism and other processes, most importantly pregnancy, in female animals. Potentially pregnant wild boar showed similar cortisol levels to non-pregnant wild boar, and there was a higher cortisol level present in pregnant wild boar, although this was only significant when potentially pregnant animals were excluded (Fig. 5). This is also seen in humans, as cortisol increases exponentially throughout pregnancy and peaks at labour [54,55]. The cortisol levels of potentially pregnant wild boar are therefore still comparable to those of non-pregnant individuals, because cortisol levels apparently increase in the last months of gestation.

To conclude, we proved differences between age and sex groups, as well as the influence of pregnancy on cortisol levels, while differences between hunting months or hunting regions could not be found. Animal welfare is becoming increasingly important in today’s society. Stress, especially long-term stress, can negatively impact the health, reproduction and longevity of wildlife, can influence the spread of diseases and can affect host-parasite equilibrium [3]. Possible chronic stress triggers include hunting methods such as drive hunting. To strengthen our findings and look further into the influence of different types of hunting on stress in wild boar, as well as possible chronic stress triggers, we need to investigate the cortisol levels of more animals under different circumstances. In addition, different media for cortisol assaying, such as saliva or faeces, should be collected throughout the year to examine basal cortisol levels, to investigate seasonal or annual changes and to explore possible chronic stress triggers. This should not only be done in wild boar, but also in other wildlife species.

## Acknowledgements

We thank all the hunters for their cooperation and help in collecting data, especially the staff from the forestry commission offices Oerrel, Unterlüß and Wolfenbüttel as well as the “Verwaltung Günther Graf v. d. Schulenburg”. Finally, yet importantly, we would like to mention the great help from Claudia Maistrelli and many volunteers and students with the fieldwork, as well as staff members of the lab with laboratory analysis. Thank you, Maura Lynch-Miller, for proofreading the language.

